# Rab35 is required for embryonic development and kidney and ureter homeostasis through regulation of epithelial cell junctions

**DOI:** 10.1101/2023.09.11.556924

**Authors:** Kelsey R. Clearman, Napassawon Timpratoom, Dharti Patel, Addison B. Rains, Courtney J. Haycraft, Mandy J. Croyle, Jeremy F. Reiter, Bradley K. Yoder

## Abstract

**Background:** Rab35 is a member of a GTPase family of endocytic trafficking proteins. Studies in cell lines have indicated that Rab35 participates in cell adhesion, polarity, cytokinesis, and primary cilia length and composition. Additionally, sea urchin Rab35 regulates actin organization and is required for gastrulation. In mice, loss of Rab35 in the CNS disrupts hippocampal development and neuronal organization. Outside of the CNS, the functions of mammalian Rab35 *in vivo* are unknown.

**Methods:** We generated and analyzed the consequences of both congenital and conditional null *Rab35* mutations in mice. Using a LacZ reporter allele, we assessed *Rab35* expression during development and postnatally. We assessed Rab35 loss in the kidney and ureter using histology, immunofluorescence microscopy, and western blotting.

**Results:** Congenital *Rab35* loss of function caused embryonic lethality: homozygous mutants arrested at E7.5 with cardiac edema. Conditional loss of Rab35, either during gestation or postnatally, caused hydronephrosis. The kidney and ureter phenotype were associated with disrupted actin cytoskeletal architecture, altered Arf6 epithelial polarity, reduced adherens junctions, loss of tight junction formation, defects in EGFR expression and localization, disrupted cell differentiation, and shortened primary cilia.

**Conclusion:** Rab35 is essential for mammalian development and the maintenance of kidney and ureter architecture. Loss of Rab35 leads to non-obstructive hydronephrosis, making the *Rab35* mutant mouse a novel mammalian model to study mechanisms underlying this disease.

**Significance Statement:** Hydronephrosis, distention of the renal calyces and pelvis, affects 1 in 100 infants. Most cases of hydronephrosis are associated with obstruction. Non-obstructive hydronephrosis is typically associated with impaired ureter development, and requires surgical intervention. Here, we describe a mouse model of non-obstructive hydronephrosis caused by mutations in *Rab35.* Hydronephrosis in *Rab35* mutants is associated with the inability to maintain epithelial cell junctions, defects in EGFR expression, and altered urothelium and smooth muscle integrity of the ureter. The *Rab35* mutant mouse is a novel model to study mechanisms and treatment strategies for non-obstructive hydronephrosis.

## Introduction

Rab35 is a member of a family of small GTPases that regulate intracellular vesicular trafficking and other cellular processes. *In vitro* studies have shown that Rab35 promotes cytokinesis through recruitment of cleavage furrow proteins [1]. Rab35 also regulates cell adhesion through recycling of E-cadherin and other adhesion proteins [2, 3]. Additionally, Rab35 antagonizes Arf6, another small GTPase that functions to internalize surface proteins, such as epithelial growth factor receptor (EGFR), that inhibit cell adhesion and promote cell migration [2]. Depletion of Rab35 in cultured kidney cells led to an inversion of cell polarity in 3D cultures, suggesting that Rab35 not only plays a role in maintaining proteins at the membrane but also their trafficking or retention in specific cellular locations [4]. In mammalian cells, Rab35 has been reported to mediate exosome secretion [5], autophagy [6–8], retrograde transport between the endosomes and Golgi [1, 2, 9], and have roles in phosphoinositide homeostasis [10–12]. Recently, Rab35 was identified as a novel regulator of primary cilia length and composition in mammalian cells and in zebrafish using siRNA- and morpholino-mediated inhibition approaches, respectively [12]. In sea urchin, Rab35 has been implicated in the regulation of actin involved in gastrulation [13]. Thus, as a small GTPase that can recruit different effectors to membranes, Rab35 may function in diverse cellular processes that is likely to be important for mammalian health and disease.

Little is known about the *in vivo* function of mammalian Rab35. Two studies demonstrated that loss of Rab35 in the CNS of mice disrupts axon elongation and distribution of hippocampal neurons, likely through altered cell adhesion [14, 15]. We found that congenital loss of Rab35 in mice is embryonic lethal. Using conditional genetics, we deleted *Rab35* late in gestation or in postnatal mice and found that either causes bilateral non-obstructive hydronephrosis, ureteral defects, and male hypogonadism. Without Rab35, kidney cells displayed mislocalization of Arf6, altered EGFR expression and localization, disrupted adherens and tight junction formation, and defects in cilia length regulation. In the ureter, there was a loss of E-cadherin expression, increased apoptosis, and diminution of smooth muscle and epithelium. These data indicate Rab35 has a critical role in maintenance of epithelial structures in the urogenital system and that Rab35 mutants represent a new mouse model for non-obstructive hydronephrosis.

## Materials and Methods

### Generation of Rab35 mutant alleles

All animal studies were conducted in compliance with the National Institutes of Health *Guide for the Care and Use of Laboratory Animals* and approved by the Institutional Animals Care and Use Committee at the University of Alabama at Birmingham. Mice were maintained on LabDiet^®^ JL Rat and Mouse/Irr 10F 5LG5 chow. The *Rab35^KO^*allele (*tm1a*) was rederived from sperm obtained from the Knockout Mouse Project (KOMP) Repository into C57BL/6J strain mice. Mice were maintained on C57BL/6J background. *Rab35* conditional allele (*tm1c*) mice were generated by mating *Rab35^KO^* to FlpO recombinase mice (C57BL/6J) to remove the LacZ and Neo cassettes generating a conditional allele (*tm1c or fl*). Progeny that contained the recombined allele were crossed off the FlpO line and bred to males carrying CAGGCre-ER (Jax Stain #004682) [16] recombinase to generate the deletion allele (*tm1d or delta*). We refer to these alleles as *tma1 (Rab35^KO^), tm1c (Rab35^fl^),* and *tm1d (Rab35^delta^)* alleles. Primers used for genotyping are included in supplemental data.

### Tamoxifen Cre Induction

Recombination of the conditional *tm1c* allele was induced in utero by injecting time pregnant females with a single intraperitoneal (IP) injection of 6 mg tamoxifen (Millipore Sigma, T5648)/40 g body weight dissolved in corn oil. Juvenile *Rab35^fl/fl^;Cre^+^*and *Rab35^fl/fl^* mice at postnatal day 7 (P7) were induced by a single IP injection of 3 mg tamoxifen/40 g body weight and adult animals at 8 weeks old by a single IP injection of 9 mg tamoxifen/40 g body weight. Cell culture media was supplemented with 1 mM 4-hydroxytamoxifen for 24 hrs to achieve deletion of Rab35 in MEFs and primary kidney epithelial cells. For simplicity throughout manuscript, all animals were induced and controls are listed as Cre- and mutants Cre+.

### Embryo Isolation

Timed pregnancies were established with embryonic timepoint of E0.5 being noon on the morning of observing the copulatory plug. To isolate embryos, pregnant females were anesthetized using isoflurane followed by cervical dislocation. Isolated embryos were transferred to PBS and yolk sac was taken for genotyping. Embryos or isolated embryonic tissues were then fixed in 4% paraformaldehyde (PFA; Sigma, 158127) in PBS.

### β-Galactosidase Staining

For whole mount or section β-galactosidase staining, samples were fixed (0.2% glutaraldehyde (Sigma), 5mM of EGTA, and 2 mM of MgCl_2_ in 1X PBS) at 4°C for 30 min, rinsed three times for 15 min at 4°C (0.02% Igepal, 0.01% sodium deoxycholate, and 2 mM MgCl_2_ in 1X PBS), and immersed in staining solution overnight in the dark at 37°C (1 mg/ml X-gal, 0.02% Igepal, 0.01% sodium deoxycholate, 5 mM potassium ferricyanide, 5 mM potassium ferrocyanide, and 2 mM MgCl_2_ in 1X PBS). Samples were postfixed in 4% PFA and stored at 4°C and sections were counterstained with Nuclear Fast Red (Sigma). Embryos and sections were imaged using a Nikon SMZ800 stereo microscope.

### Mouse embryonic fibroblast (MEF) Isolation

Timed pregnancies between *Rab35^fl/fl^* and *Rab35^fl/fl^;Cre+* animals were set up to isolate embryos at E13.5 to culture mouse embryonic fibroblasts (MEFs). Following the removal of the liver and head, embryos were mechanically dissociated and cultured in DMEM (Gibco, 11039-021) supplemented with 10% Fetal Bovine Serum, 1X Penicillin and Streptomycin, 0.05% Primocin, 3.6μl/0.5L β-mercaptoethanol. MEFs were cultured on glass cover slips treated with 0.1% gelatin 24-48 hrs prior to tamoxifen induction. To induce cilia formation, cells were grown to confluency and then switched to DMEM medium containing 0.5% FBS [17].

### Primary kidney epithelial cell isolation

1-month-old *Rab35^fl/fl^;Cre+* animals were anesthetized with isoflurane followed by cervical dislocation and kidneys were removed and mechanically dissociated in a sterile environment. Minced tissue was filtered through a 70-µm cell strainer and resuspended in DMEM (Gibco, 11039-021) supplemented with 5% FBS, epidermal growth factor (recombinant human, 10ng/ml), insulin (recombinant human, 5 µg/ml), hydrocortisone (36 ng/ml), epinephrine (0.5 µg/ml), Triiodo-L-thyronine (4 pg/ml), and transferrin (recombinant human, 5 µg/ml) (Growth Medium 2 Supplement Pack, PromoCell, C-39605). Cells were allowed to grow until 80-90% confluent and culture medium promotes epithelial cell survival. Primary epithelial cells were cultured 48-72 hrs prior to tamoxifen induction. Primary kidney epithelial cells were cultured for a maximum of 4 passages.

### Tissue Isolation and Histology

Mice were anesthetized with 0.1 ml/10 g of body weight dose of 2.0% tribromoethanol (Sigma Aldrich, St. Louis, MO) and transcardially perfused with PBS followed by 4% PFA. Tissues were post-fixed in 4% PFA overnight at 4°C and then cryoprotected by submersion in 30% sucrose in PBS for 24 hours then frozen in O.C.T. (Fisher Scientific, 23-730-571) and cryosectioned at 10 µm or 20 µm thickness for immunofluorescence or hematoxylin (Fisher Chemical, SH26-500D) and eosin (Sigma-Aldrich, HT110132-1L) staining was performed.

### Necropsy Analysis

Animals were sent to the Comparative Pathology Lab (UAB) for necropsy where all tissues were fixed in 10% neutral buffered formalin overnight, processed into 5 µm sections, and stained with hematoxylin and eosin. Slides were evaluated for tissue histopathology by a board-certified veterinary pathologist. For blood chemistry analysis, blood was collected by axial bleed and provided to the Comparative Pathology Lab to measure serum calcium, blood urea nitrogen (BUN), creatinine, albumin, sodium, phosphate, and total protein. All analyses were performed blinded.

### Immunofluorescence Microscopy

Tissue sections (10 µm, except kidney sections used for cilia length measurements which were 20 µm) were fixed in 4% PFA for 10 minutes, permeabilized with 0.1% TritonX-100 in PBS for 8 minutes and then blocked in a PBS or TBS (depending on antibodies) containing 1% BSA, 0.3% TritonX-100, 2% normal donkey serum and 0.02% sodium azide for one hour at room temperature. Primary antibodies were incubated in blocking solution overnight at 4°C. Detailed information about primary antibody usage is in **Supplemental Data 1**. Cryosections were then washed with PBS/TBS three times for five minutes at room temperature. Secondary antibodies diluted in blocking solution were added for one hour at room temperature. Secondary antibodies were donkey conjugated Alexa Fluor 647, 594 or 488 (Invitrogen, 1:1000). Samples were then washed in PBS and stained with Hoechst nuclear stain 33258 (Sigma-Aldrich) for 5 minutes at room temperature. Cover slips were mounted using Immu-Mount (Thermo Scientific). All fluorescence images were captured on Nikon Spinning-disk confocal microscope with Yokogawa X1 disk, using Hamamatsu flash4 sCMOS camera. We used 60x apo-TIRF (NA=1.49), 40x plan fluor (NA=1.3) or 20x Plan Flour multi-immersion (NA=0.8) objectives. Images were processed using Nikon Elements or Fiji software.

### Intrapelvic Dye Injections

Timed pregnant females were injected with tamoxifen at E14.5 and euthanized at E16.5 for embryo isolations. Yolk sacs were taken for genotyping and dye injection was done blinded prior to genotyping. Whole kidneys with ureters and bladder attached were removed from embryos and a 0.1% solution of Bromo-Phenol Blue in PBS was injected into the kidney pelvis through a pulled glass pipette. Injections were recorded by timelapse imaging and then converted into movie files. Ureters were recorded post dye-injection for 5 minutes by timelapse imaging.

### Statistical Analysis

Analyses were performed using GraphPad Prism and Microsoft Excel. Specific tests used are indicated in figure legends with significance indicated as follows: * p≤0.05, ** p≤0.01, *** p≤0.001. All error bars represent standard error of the mean (SEM).

## Results

### Rab35 expression in the mouse

The *Rab35 tm1a* allele (*Rab35^KO^*) contains a LacZ reporter encoding β-galactosidase (**Figure 1a**). β-galactosidase staining of E8.5 *Rab35 tm1a* heterozygous embryos revealed that *Rab35* was ubiquitously expressed (**Figure 1b**). At P7, β-galactosidase activity was more restricted to the kidney, ureter, and testes (**Figure 1c-e**). Modest β-galactosidase activity was present in the lung and liver, and no β-galactosidase activity was evident in the heart and pancreas (**Figure 1f-i**). In tissues where β-galactosidase activity was detected, it was enriched in epithelial cells and ciliated cells (e.g., biliary epithelial cells and spermatozoa, **Figure 1g**). In the kidney, β-galactosidase activity was not uniform, with greater activity in the cortex and renal papilla than in the medulla (**Figure 1c-c’**). β-galactosidase activity was also not uniform in the ureter, with greater activity in the urothelium than in the underlying smooth muscle (**Figure 1d-d’**).

**Figure 1.**
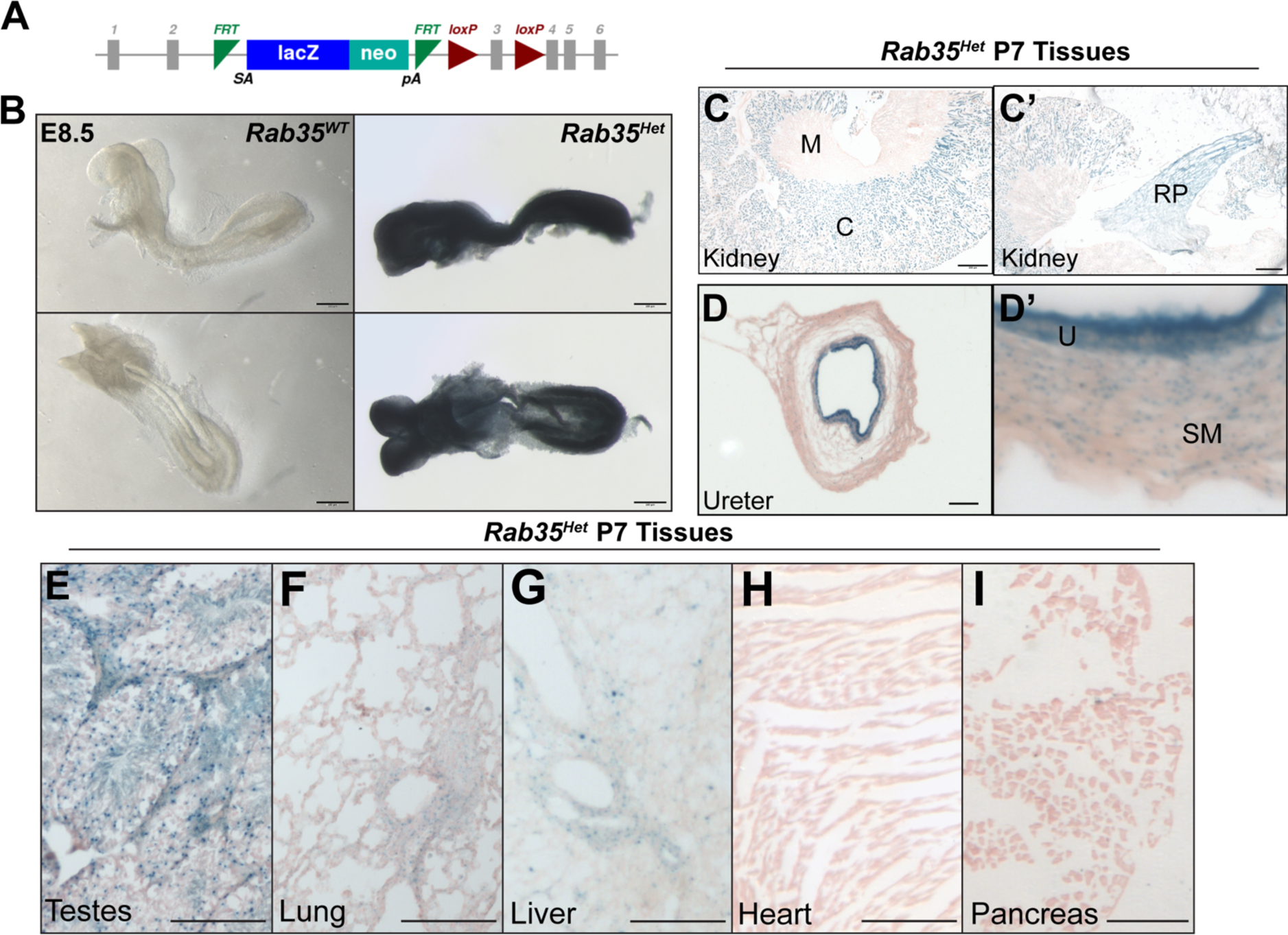
*Rab35* expression in the mouse. **A)** The *tma1* allele that was used to generate *Rab35^KO^*mice contains a LacZ reporter. β-galactosidase staining revealed *Rab35* expression in embryos and postnatal tissues. **B)** Staining for β-galactosidase activity in E8.5 heterozygous embryos indicated that *Rab35* is ubiquitously expressed. β-galactosidase activity in P7 tissues was more restricted to **C)** kidney (M, medulla, C, cortex, and RP, renal papilla) and **D)** ureters (U, urothelium, SM, smooth muscle). β-galactosidase activity was also present in **E)** testes**, F)** lung, **G)** liver, but not in **H)** heart and **I)** pancreas.

### Rab35 regulates primary cilia length, but is not required for ciliogenesis in mice

Previous studies have shown that Rab35 depletion using siRNA in mammalian cells or morpholinos in zebrafish reduces primary cilia length [12]. To confirm whether Rab35 regulates cilia length in mammalian cells, we generated primary mouse embryonic fibroblasts (MEFs) and renal epithelium from conditional *Rab35^tm1c/tm1c^;CAGG-CreER* mice (**Supplemental Figure 1a**). *Rab35* deletion was induced by addition of 4-hydroxytamoxifen to the culture medium to generate *Rab35^tm1d/tm1d^;CAGG-CreER* cells (hereafter referred to as *Cre+* cells). The percentage of cells with cilia and cilia length was quantified after staining for acetylated α-tubulin, a ciliary component. The percentage of *Cre+* cells with a cilium was not significantly different than the percentage of control cells with a cilium. However, MEFs and epithelial cells lacking Rab35 had shorter cilia (**Supplemental Figure 1b-g**). Similarly, deletion of *Rab35 in vivo* reduced primary cilia length in the kidney or ureter (**Supplemental Figure 1h-k**). Therefore, mammalian Rab35 maintains cilia length and elongation, but is not essential for ciliogenesis.

### Rab35 is essential for embryonic development

No homozygous *Rab35^KO^* embryos were isolated after E10.0, indicating that Rab35 is required for embryonic development (**Figure 2a-b**). *Rab35^KO^* embryos isolated at E8.5 were developmentally delayed (**Figure 2c**). *Rab35^KO^* embryos had disorganized filamentous actin (F-actin, **Figure 2d**). As Rab35 is required for actin organization in sea urchins [13], Rab35 function is evolutionarily conserved.

**Figure 2.**
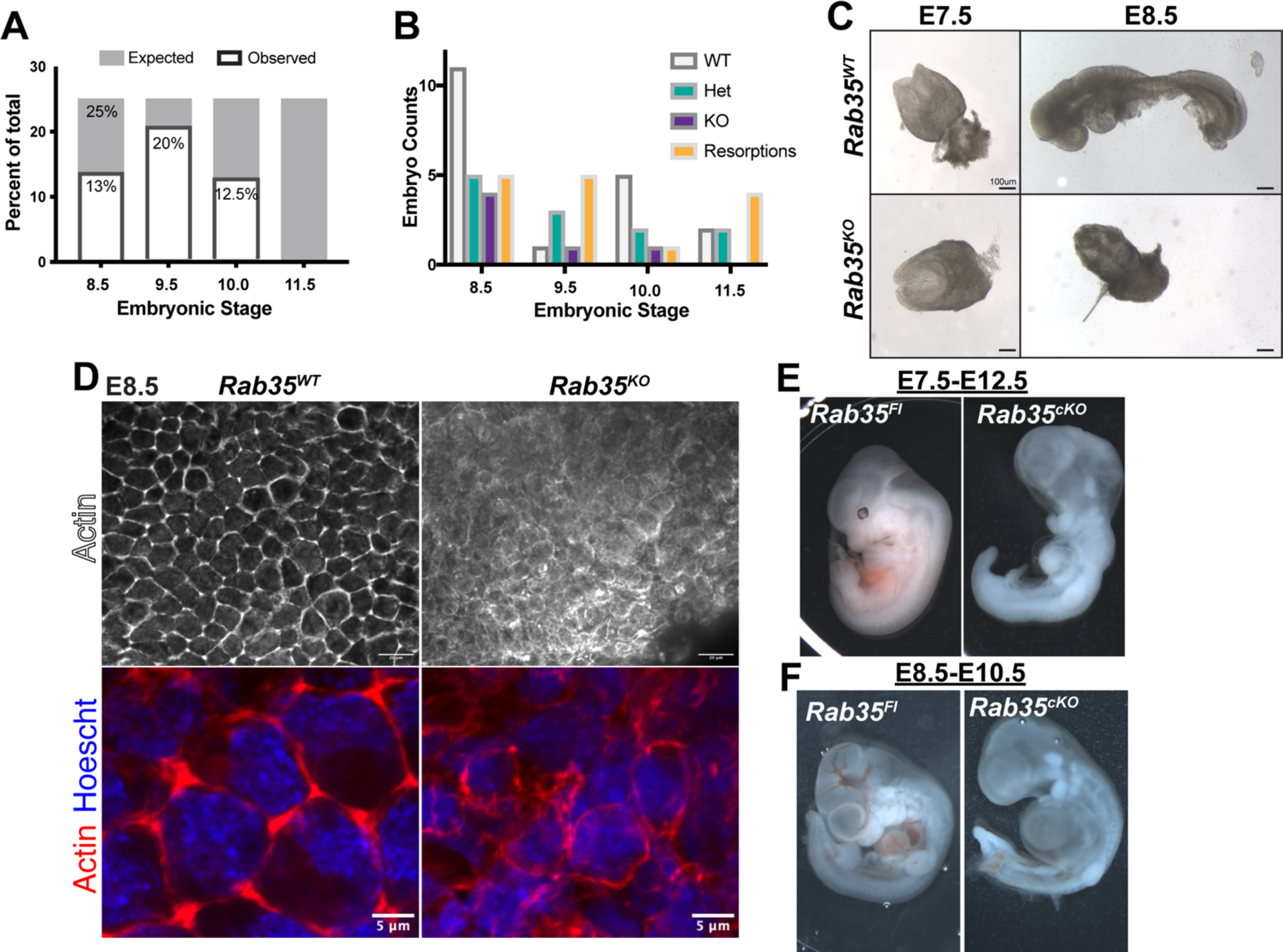
Rab35 is required for embryonic development in the mouse. **A)** Embryos were isolated between E8.5 and E11.5 and genotyped. Until E10.0, *Rab35^KO^* mutant embryos were observed at sub-Mendelian ratios. *Rab35^KO^*mutant embryos were not identified at E11.5. **B)** Quantitation of embryo genotypes between E8.5 and E11.5. **C)** E7.5 *Rab35^KO^* embryos were morphologically comparable to wild-type sibling controls. At E8.5, *Rab35^KO^*embryos were developmentally delayed. **D)** Phalloidin staining of wild-type sibling controls and *Rab35^KO^* embryos **E)** Conditional deletion of *Rab35* at E7.5 and analysis at E12.5 and **F)** deletion at E8.5 with analysis at E10.5 reveals cardiac edema and developmental delay.

### Conditional deletion of Rab35 causes bilateral hydronephrosis and decreased kidney function

To circumvent the early lethality in *Rab35* germline null mutants and uncover other functions of Rab35, we deleted *Rab35* during gestation (**Figure 2e-f**, **Figure 3a-b**) using a ubiquitously expressed tamoxifen-inducible Cre transgene (CAGG-CreER, Cre+) (**Supplemental Figure 1a**). Deletion of *Rab35* at E7.5, prior to organogenesis, resulted in developmental arrest and lethality (**Figure 2e**). We also induced *Rab35* deletion at E8.5 and analyzed embryos at E10.5. Although Kuhns et al implicated Rab35 in regulating ciliary Shh signaling, we did not detect phenotypes typically associated with altered Hedgehog signaling (e.g., polydactyly)[12]. Deletion of Rab35 at E8.5 resulted in developmental arrest, cardiac edema, and lethality (**Figure 2f**).

**Figure 3.**
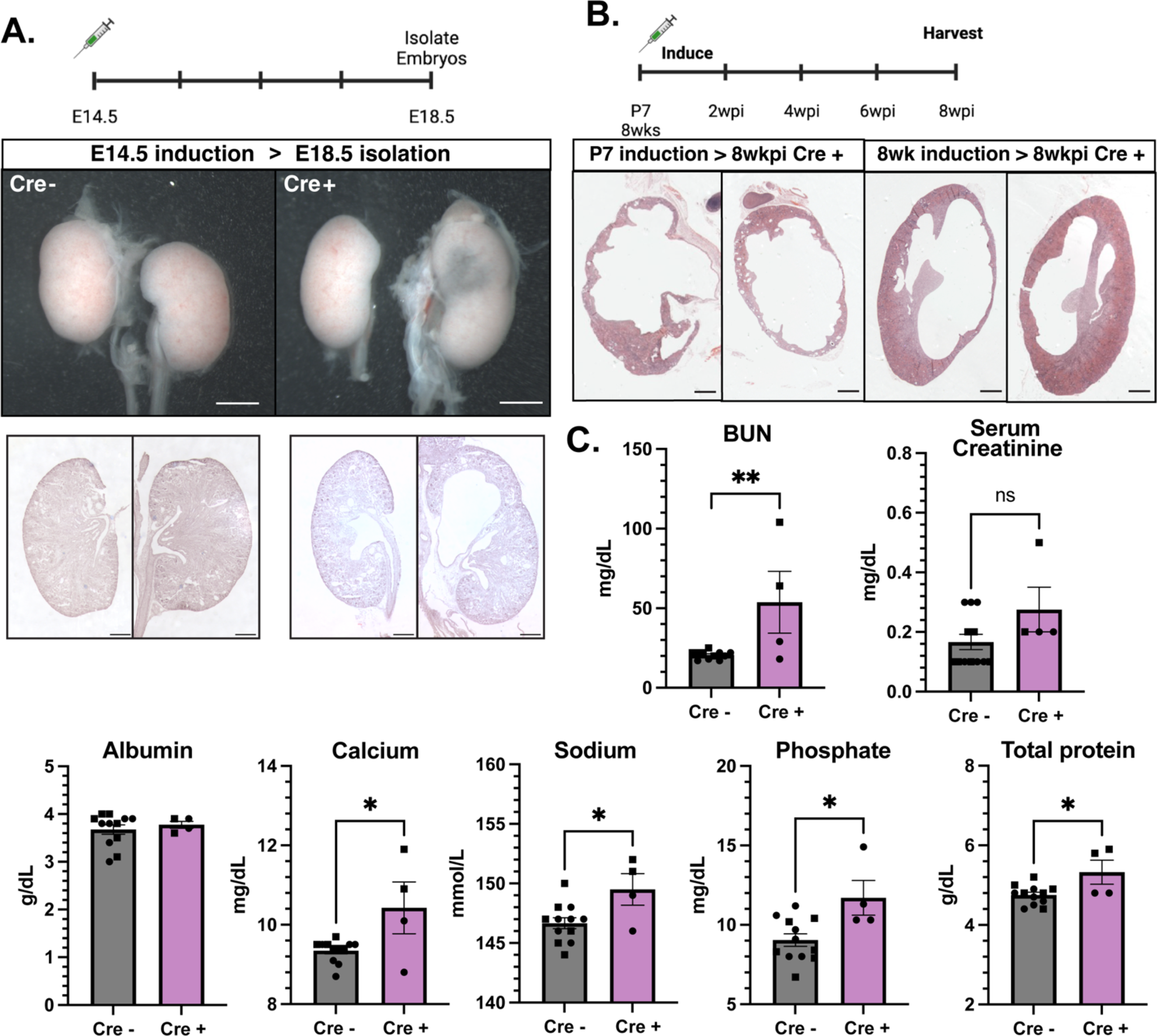
Conditional deletion of Rab35 leads to bilateral hydronephrosis and decline in kidney function. **A)** Timeline for Rab35 embryonic inductions. *Rab35^delta^;Cre+* embryos isolated at E18.5 had bilateral hydronephrosis observed by gross morphology and histological staining of 10µm sections. **(B)** Timeline for Rab35 juvenile (P7) and adult (8wk) inductions. Independent of induction time, *Rab35^delta^;cre+* mice have bilateral hydronephrosis 8 weeks post induction (wkpi). **(C)** Blood serum analysis of kidney function markers. Blood urea nitrogen (BUN) P value= 0.0098, calcium P value=0.0130, sodium P value= 0.0208, phosphate P value= 0.0113, total protein P value= 0.0139.

Deletion of *Rab35* at E14.5, mid-organogenesis, and analysis of embryos at E18.5 revealed severe bilateral hydronephrosis in *Cre+* embryos (**Figure 3a**). To determine if the hydronephrosis was necessarily of developmental origin, we deleted *Rab35* postnatally in juvenile (P7) and adult (8wk) conditional mice (**Figure 3b**). All *Cre+* animals induced at either P7 or at 8wk and analyzed 8 weeks post-tamoxifen induction (wkpi) exhibited hydronephrosis, indicating that Rab35 is required postnatally to maintain renal architecture. Blood serum chemistry analysis of the *Cre+* animals induced at P7 revealed increased blood urea nitrogen (BUN), calcium, sodium, phosphate, and total protein, indicative of decreased renal function (**Figure 3c).** There was no significant difference in serum creatinine and albumin in *Cre+* animals compared to littermate controls at 8 wkpi (**Figure 3c**).

To characterize the onset of hydronephrosis in *Cre+* embryos and in postnatal mice, we assessed kidneys at a series of time points after Rab35 deletion at E14.5 or P7 (**Supplemental Figure 2**). Following deletion at E14.5, E16.5 *Cre+* embryos did not display hydronephrosis (**Supplemental Figure 2a**). Similarly, following deletion at P7, 4 wpki *Cre+* mice did not display hydronephrosis (**Supplemental Figure 2b**), but did at 6 wpki. To assess the pathogenesis of the hydronephrosis caused by Rab35 deletion, we therefore examined E16.5 *Cre+* embryos and 4 wpki *Cre+* mice.

### Rab35 participates in renal epithelial cell adhesion and polarity

*In vitro* studies indicate that Rab35 is important for maintaining epithelial cell junctions through opposing the activity of Arf6 and through E-cadherin membrane recycling [2, 3]. Analysis of Arf6 localization in adult and embryonic *Cre+* kidney tubule epithelium showed an expansion of Arf6 distribution across the cell membrane, instead of its normal restriction to the apical epithelium (**Figure 4a and Figure 5a**). This expanded distribution was observed prior to the onset of hydronephrosis. Western blot analysis revealed that the expanded distribution could not be explained by increased Arf6 protein (**Figure 4b**).

**Figure 4.**
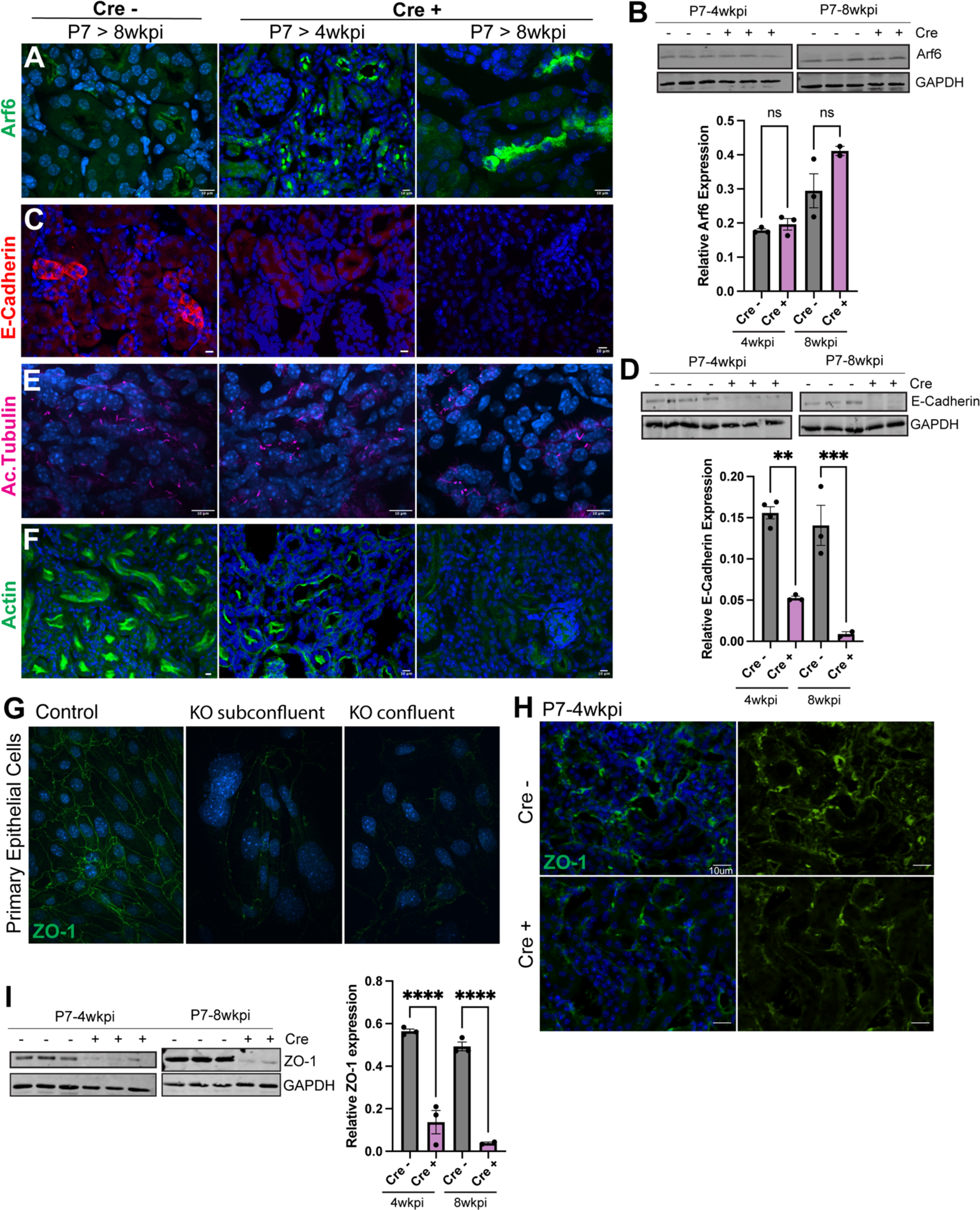
Rab35 regulates epithelial cell adhesion and polarity within the adult kidney. **A)** IF staining for Arf6 in control, pre-hydronephrotic (4wkpi) and hydronephrotic (8wkpi) kidneys. **B)** Western blot of Arf6 in kidney lysates from control, pre-hydronephrotic, and hydronephrotic and quantification normalized to GAPDH. **C)** IF staining for E-cadherin and **D)** western blot for E-cadherin with quantification in whole kidney lysates normalized to GAPDH. **E)** Alpha-acetylated tubulin staining indicated no change in primary cilia polarity in kidneys. **F)** Phallodin shows a decrease in actin organization and brush border. **G)** Tight junction marker, ZO-1 is reduced in *Rab35^delta^;Cre+* cultured primary epithelial cells independent of confluency and **H-I)** significantly reduced in pre and post hydronephrotic kidneys. Statistical analysis was done using one-way ANOVA and post hoc analysis for multiple comparisons between each timepoint. Pre-hydronephrosis analysis was not compared to hydronephrosis analysis. P value for 4wkpi (pre-hydronephrosis) quantification of E-cad is 0.0011 and for 8wkpi (hydronephrosis) analysis, P value = 0.0007. P values for ZO-1 <0.0001.

**Figure 5.**
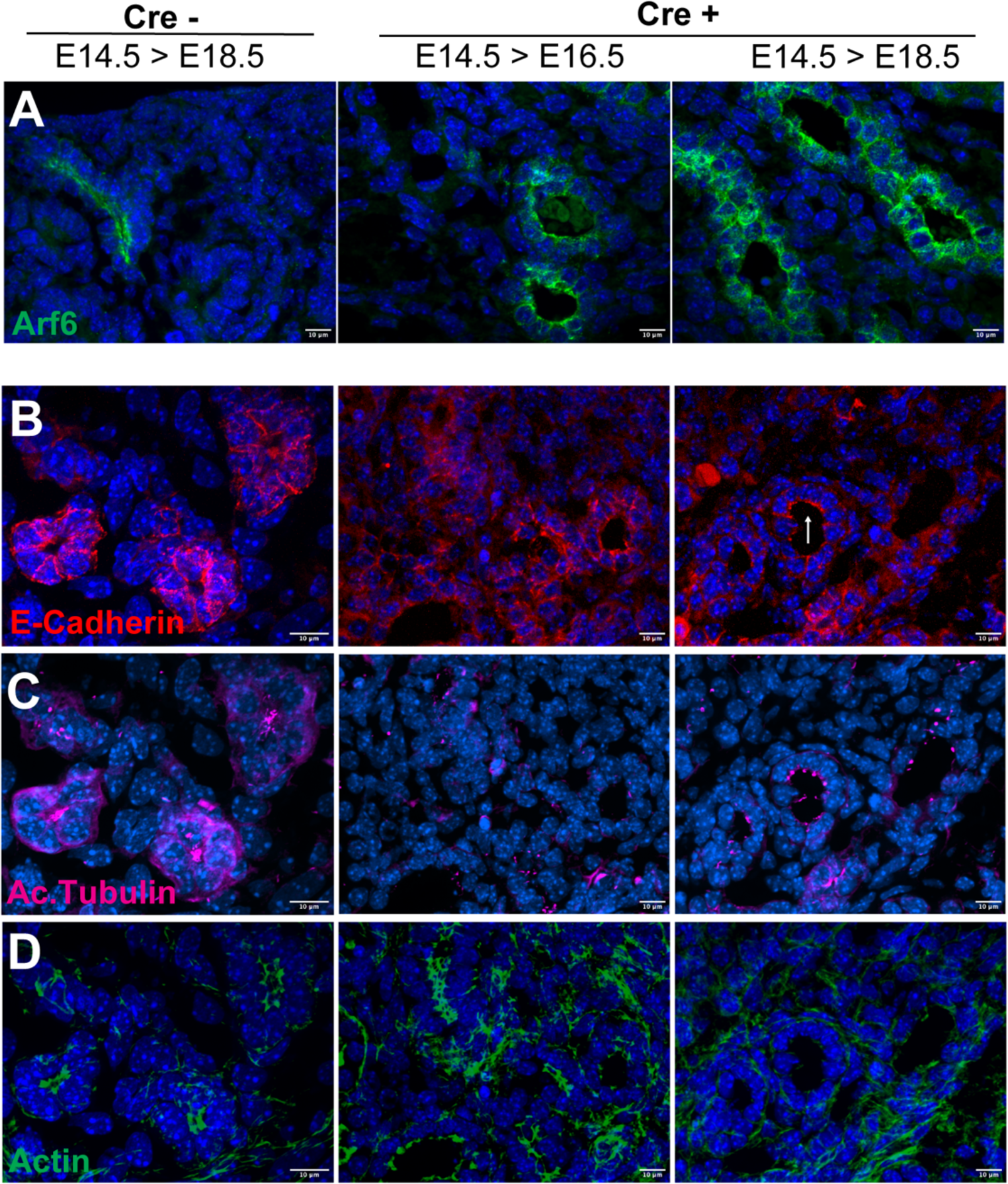
Rab35 regulates epithelial cell adhesion and polarity within the embryonic kidney. **A)** IF staining for Arf6 in control, pre-hydronephrotic (E16.5), and hydronephrotic (E18.5) kidneys shows similar localization changes as in adult kidneys. **B)** E-cadherin is reduced and has changes in localization from baso-lateral to apical membranes (white arrow). **C)** Primary cilia are present and apically oriented. **D)** Reduction in brush border actin.

Along with expanded Arf6 localization, *Cre+* kidney tubule epithelium had reduced E-cadherin and *Cre +* embryonic kidney tubule epithelium had apical E-cadherin expression (**Figure 4c and Figure 5b**). Quantification of E-cadherin by western blot analysis showed reduced E-cadherin levels prior to the onset of hydronephrosis (**Figure 4d**). In addition to E-cadherin, *Cre+* kidneys displayed a reduction in N-cadherin (**Supplemental Figure 3**), although to a lesser degree and its localization was not altered. Following this same trend, primary cilia, although shorter, remained apically oriented in *Cre+* kidneys (**Figure 4e and Figure 5c**).

With the increased membrane localization of Arf6 and reduced E-cadherin maintaining adherens junctions, we suspected an overall reduction in epithelial cell junction integrity and adhesion. To assess this, we analyzed the tight junction marker, ZO-1 by IF and western blot using *Cre+* primary epithelial cells and *in vivo* in the kidney (**Figure 4g-i**). In both, there is a significant reduction of ZO-1 evident pre- and post-hydronephrosis (**Figure 4i**). These data support a model in which Rab35 is required to maintain E-cadherin mediated epithelial cell adherens and ZO-1 at tight junctions, within the kidney. It is likely that the effect of Rab35’s loss on tight junctions is indirect as adherens junctions are required for tight junction formation and maintenance [18].

Along with defects in maintaining junctional complexes in *Cre+* kidneys, there is a loss of actin organization and more specifically, a loss of brush border actin (**Figure 4f and Figure 5d**) that may contribute to a loss of polarity and epithelial junction integrity. To assess polarity defects, we analyzed EGFR localization, which is normally restricted to the basal-lateral membrane of kidney epithelia. Further, data from other groups indicate that EGFR localization is regulated by E-cadherin and Rab35/Arf6 activity *in vitro* [19, 20]. In *Cre+* kidneys, EGFR is no longer restricted to the basal-lateral membrane pre- or post-hydronephrosis and its expression is significantly increased in hydronephrotic kidneys (**Figure 6a-c**). This suggests that Rab35 regulates EGFR internalization and degradation *in vivo* in the kidney in addition to regulation of E-cadherin and Arf6 membrane localization and polarity. However, our analysis *in vivo* indicates there is not a complete loss of epithelial cell polarity or organization in Rab35 mutant cells as CD13 remains apically localized (**Figure 6a**) as does the position of the primary cilium (**Figure 4e and Figure 5c**).

**Figure 6.**
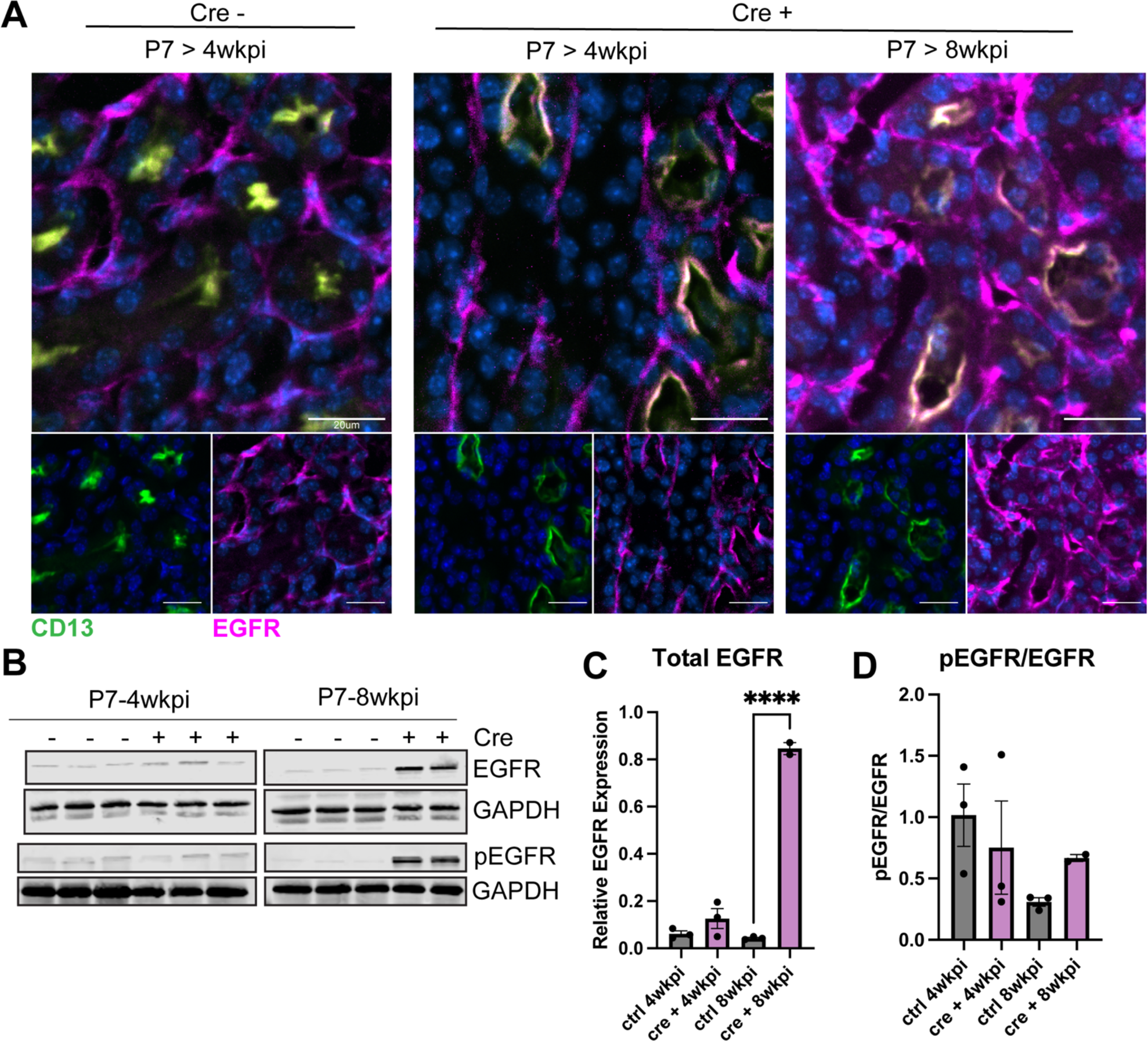
Rab35 regulates EGFR expression and localization within the kidney. **A)** IF of EGFR expression in control, pre-hydronephrotic, and hydronephrotic kidneys. CD13 is used as a marker for apical membrane in kidney epithelia to show changes in EGFR localization in *Rab35^delta^; Cre+* kidneys. **B)** Western blot of whole kidney lysates and **C)** quantification of EGFR normalized to GADPH. Only hydronephrotic kidneys had significant increases in EGFR with a P value <0.0001 **D)** A ratio was taken of phosphorylated EGFR out of total EGFR to determine if there were changes in EGFR activity. There was no significant difference in pEGFR/EGFR ratio across kidney samples.

### Rab35 regulates ureter integrity through maintenance of the urothelium

Bilateral hydronephrosis is most commonly caused by an obstruction along the kidney-ureter-bladder tract. However, in *Cre+* animals there is no evidence of obstruction, as adult animals urinate normally and dye injections in *Cre+* embryos show the lumen of the ureter remains patent (**Figure 7a-b and Supplemental Video 1-2**). Thus, Rab35 mutants are a new model of non-obstructive hydronephrosis. Interestingly, the lumens of *Cre+* mutants were dilated compared to the *Cre-* control ureters (**Supplemental Video 2, Figure 7a-b**). Histological analysis of cross sections through the ureters revealed the *Cre+* ureters were thin even at the pre-hydronephrotic stage (**Figure 7c**).

**Figure 7.**
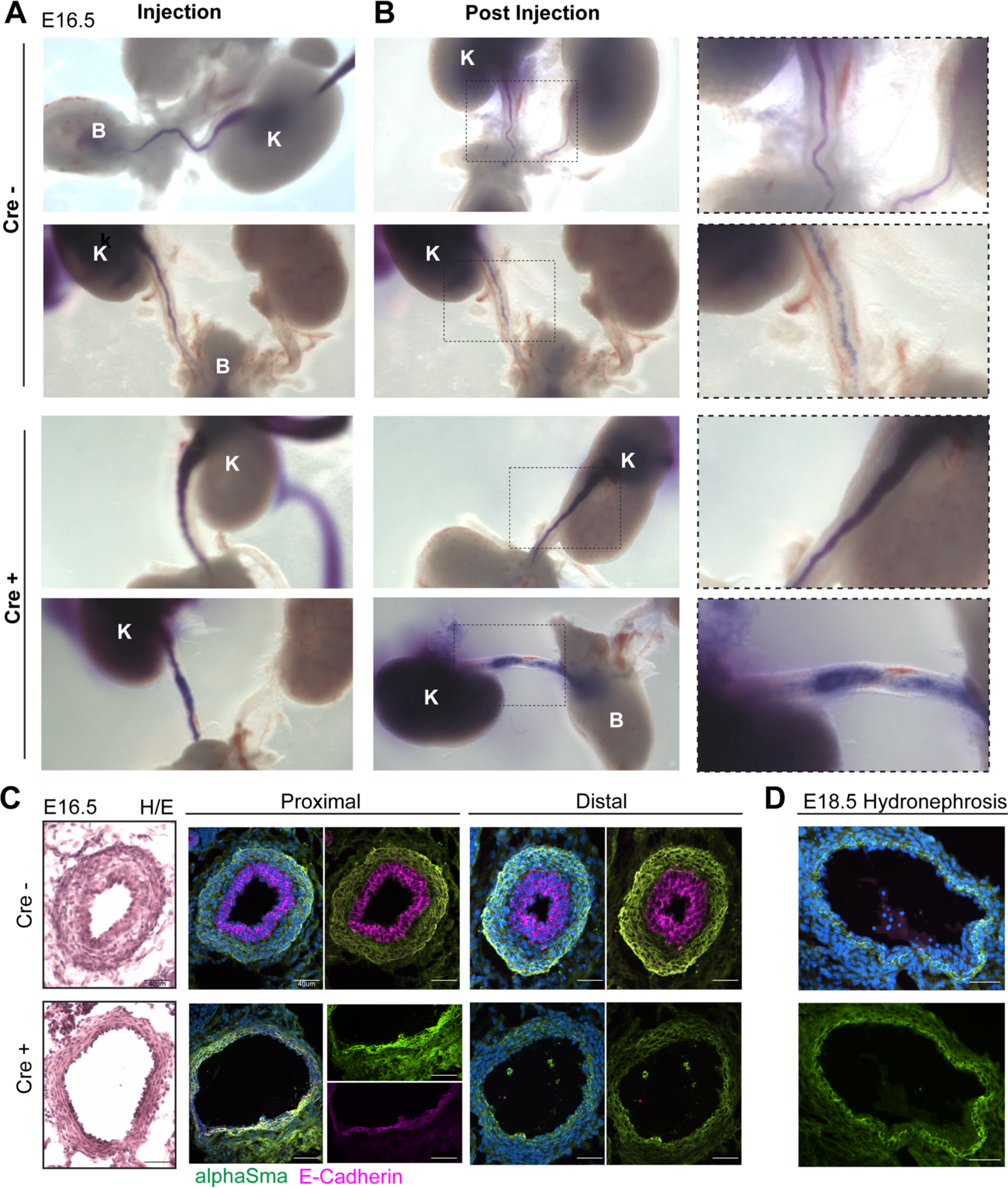
Rab35 regulates ureter epithelium and smooth muscle maintenance. Embryonic intrapelvic dye injections suggest defects in ureter lumen structure. **A)** Snapshot of dye injection in two *Rab35^fl^* kidneys and two *Rab35^delta^;Cre+* kidneys (K, Kidney and B, Bladder) **B)** 3-minute snapshot post dye injection suggest widening of lumens. **C)** Histological analysis of pre-hydronephrotic embryonic ureters by H and E staining confirmed widening of ureter lumens in *Rab35^delta^;Cre+* embryos. IF staining of E-cadherin and smooth muscle actin, showed *Rab35^delta^;Cre+* proximal ureters had cells expressing both epithelial and smooth muscle markers where the distal ureters no longer had any E-cadherin expressing cells. This was similar to **D)** ureters isolated after hydronephrosis (E18.5).

Since loss of Rab35 led to a reduction in E-cadherin in the kidney, we analyzed whether there is a similar reduction in the urothelium associated with the widening of the ureter lumen. The data confirm a marked reduction in E-cadherin expression in the proximal regions of the ureter and that the distal regions of the ureter lack E-cadherin positive urothelium (**Figure 7c-d**). Normally the urothelium (E-Cad+) and smooth muscle (aSMA+) cells are localized in well-defined regions of the ureter. However, cells in the *Cre+* proximal ureters co-stained for both smooth muscle actin and E-cadherin, suggesting these cells had lost their normal identity (**Figure 7d**). No overt defects in smooth muscle are evident in the kidney, and there were no other indications that smooth muscle in general was impaired in *Cre+* as indicated by necropsy and IF staining in other tissues (**Supplemental Figure 4**).

Since deletion of Rab35 was induced after the ureter epithelium and smooth muscle layers had been established, we hypothesized that the thinning of the ureter was due to loss of these cells through apoptosis. In the pre-hydronephrotic *Rab3^delta^;cre+* kidney and ureter there is an increase in expression of cleaved-caspase-3 (CC3) supporting this hypothesis. This is specific to the epithelial cells that no longer expressed E-cadherin and was observed prior to the onset of hydronephrosis (**Figure 8a-b**).

**Figure 8.**
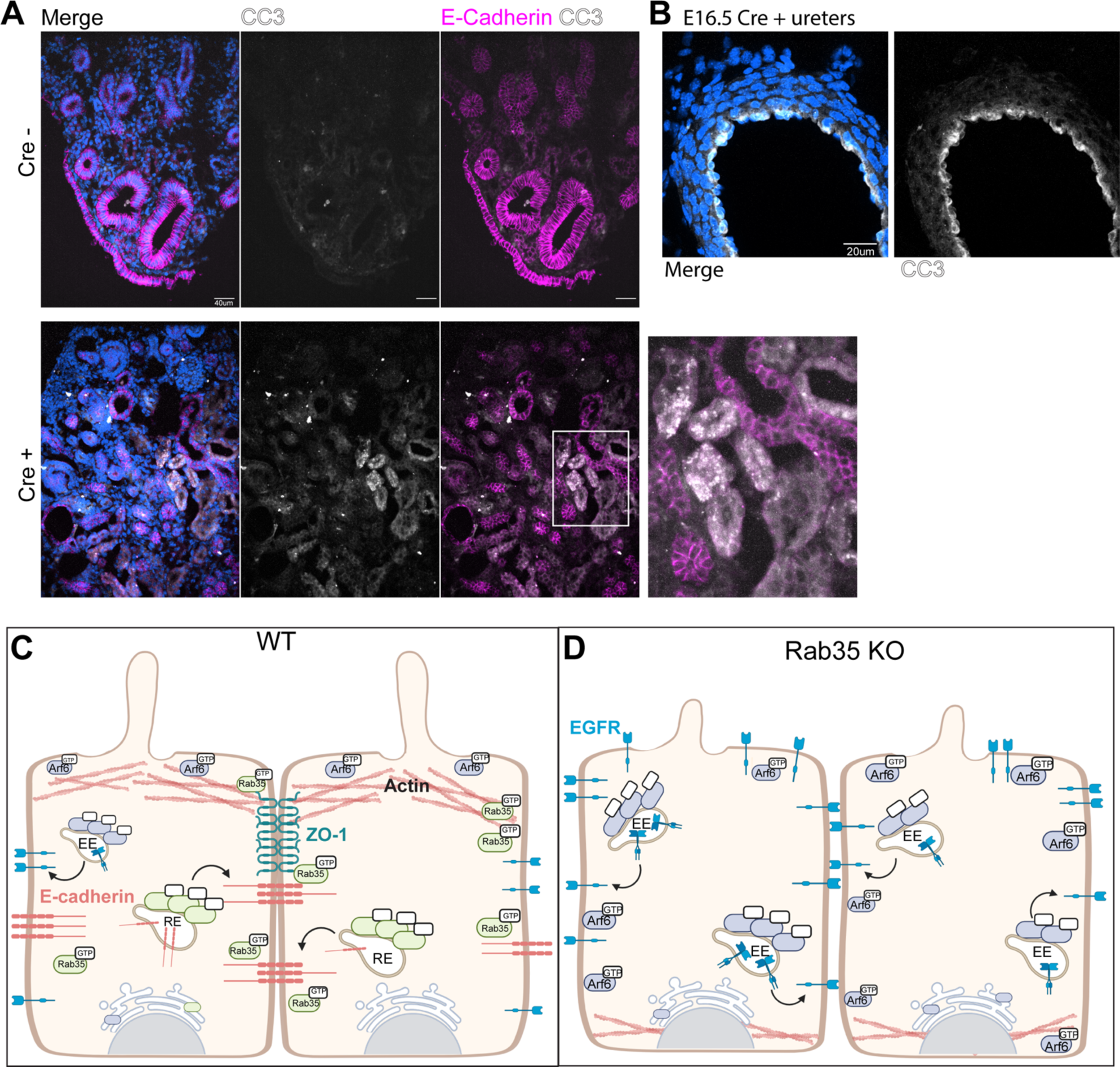
Loss of Rab35 leads to apoptosis in the kidney and ureter. **A)** IF staining of kidneys with E-cadherin and a marker for apoptosis, cleaved-caspase 3, indicated cell death was occurring in cells no longer expressing E-cadherin. This was also observed in **B)** ureters **C)** Working model: Rab35 maintains actin organization and E-cadherin membrane recycling in addition to inhibiting Arf6 activity. This creates a spatial boundary for Arf6 that is lost when Rab35 is removed. EGFR membrane expression is finely tuned by the presence of active Rab35 and E-cadherin. Arf6 activation leads to the internalization and recycling of EGFR to the membrane. Loss of these boundaries set by Rab35 through E-cadherin leads to increased recycling of EGFR, reduced adherens junctions, and loss of tight junction protein ZO-1. These changes in epithelial cells within the ureter and kidney lead to loss of epithelial integrity, cell death, and hydronephrosis.

Collectively, these data suggests that Rab35 is required for the maintenance of the urothelium through regulation of E-cadherin and loss of this regulation subsequently affects the surrounding smooth muscle cells. Interestingly, this regulation is specific to the ureter and kidney as other tissues in *Cre+* embryos and adults that we analyzed have E-cadherin and smooth muscle actin expression comparable to *Cre-* at these timepoints (**Supplemental Figure 4**). We suspect that this may be due to the time point analyzed with the kidney being more sensitive to Rab35 loss than other tissues, and thus maybe the first tissue in which a phenotype manifests.

## Discussion

### Rab35 as a novel regulator of epithelial cell adhesion and polarity in kidney development and maintenance

Rab35 is a critical regulator of many cellular processes including cilia length, vesicle trafficking, cytoskeletal organization, cell adhesion and migration, cytokinesis, and axonal elongation. While loss of Rab35 is essential for viability, very little is known in mammals regarding the impact of Rab35 functional loss in tissues other than in the central nervous system. Using a congenital *Rab35^KO^* reporter allele, we showed that Rab35 is ubiquitously expressed in early gestational embryos but has more restricted expression in the postnatal mouse with highest expression in the kidney and ureter. We have shown that Rab35 congenital null embryos die prior to E8.5, while others have reported viable embryos at E10.5 [15].

We then assessed whether mutant embryos are viable when loss is induced at later developmental timepoints. To address this, we utilized conditional null allele of Rab35 to bypass the early gestational lethality and evaluated the importance of Rab35 in late gestation and in postnatal tissues. Although we noted a decrease in cilia length in the absence of Rab35, our analysis of early embryos did not show any overt signs of left-right axis or neural tube closure defects, both classic ciliopathy phenotypes. This indicated it is not sufficient to cause cilia functional loss although this may be confounded by the development arrest in the mutants. A consistent phenotype that we do observe in Rab35 mutants, and a likely contributor to their lethality, is a marked alteration in the actin cytoskeletal organization. This phenotype was also reported in sea urchins lacking Rab35 indicating conserved function [13]. Rab35 is known to recruit effectors that modulate localized actin assembly and organization [21], including Arf6 [2, 10, 11], RhoA and Rac1 [22–24], in multiple cell types. This regulation is critical for many cell processes that are important for normal development.

To evaluate Rab35 mutant perinatal and postnatal phenotypes, we induced Rab35 deletion at E14.5 and P7 after organogenesis has occurred. In both cases, Rab35 mutants have severe and fully penetrant bilateral hydronephrosis. Hydronephrosis is not considered widely as a ciliopathy, although defects in the hedgehog pathway in the smooth muscle surrounding the ureter can cause this phenotype [25, 26]. However, we did not observe typical ciliopathy renal phenotypes such as cyst formation.

As we observed in the Rab35 mutant embryos, there is changes in actin cytoskeletal organization from its apical enrichment in kidney epithelial cells to a diffuse pattern lacking organization in *Cre+* kidneys. F-actin can regulate epithelial adherens junctions, polarity, and organization, all of which Rab35 has been shown to regulate *in vitro*. To determine the molecular/cellular mechanism involved in hydronephrosis, we investigated Arf6 and E-cadherin expression and localization, which have been shown to be affected in previous studies. Interestingly, we observed a redistribution of Arf6 from an apical domain to both apical and basal lateral membranes in the Rab35 mutants. This is indicative of an overactivated Arf6 [27, 28]. We also found a lack of E-cadherin at the cell junctions and a loss of E-cadherin protein expression that would disrupt the adherens junctions. As Arf6 and Rab35 are counter pathways with Arf6 promoting membrane protein internalization, this could explain the large decrease in E-cadherin. The loss of Rab35 does not only affect the adherens junctions as we observe a similar effect on ZO-1. Tight junction proteins have not been previously shown to be regulated by Rab35 and this may be a consequence of changes within the actin cytoskeleton and loss of E-cadherin at adherens junctions [29]. The effects on the adherens and tight junctions occur prior to the onset of hydronephrosis and are present both *in vivo* and in primary renal cells cultured from the Rab35 mutants. Thus, disruption of the adherens and tight junctions is not a secondary consequence of hydronephrosis. Interestingly, the changes in polarity and expression of these adherens and tight junction proteins were specific to Arf6, E-cadherin, and ZO-1, as N-cadherin remained localized to basal-lateral membranes despite there being data indicating its regulation by Rab35 in cell culture [3]. Additionally, apical/basal polarity is not fully disrupted as primary cilia and CD13 remain in apical orientation, although cilia are shorter than normal.

The extracellular domain of E-cadherin binds to EGFR and this interaction helps create the boundary for EGFR localization while in turn EGFR activity and signaling can regulate E-cadherin internalization and expression [19, 29–32]. Arf6 has also been shown to internalize E-cadherin in response to EGFR activity and this internalization requires Arf6 activity [33]. While at later stages, we observe an increase in EGFR expression and changes in localization, it is only seen after significant changes in Arf6 and E-cadherin are present suggesting the changes in EGFR are a consequence of the loss of E-cadherin and ZO-1 and increased Arf6 localization. This data further suggests that Rab35 may be involved in EGFR internalization and degradation in the kidney through regulation of these pathways. Previous studies have identified Rab35 to be a regulator of EGFR expression and activity is various human cancer cell lines and this regulation is dependent upon Rab35 activity [2, 20, 34, 35].

### Rab35 as a novel regulator of ureter organization

As we found no obstruction in our model and Rab35 is highly expressed in the urothelium and underlying smooth muscle, we hypothesized that there would be similar epithelial changes in the ureter as in the kidney. Like in *Rab3^delta^;cre+* kidneys, there was a dramatic reduction in E-cadherin positive cells, and those that remained E-cadherin positive also expressed smooth muscle actin. This suggests that Rab35 is required to maintain the epithelial cell integrity within the urothelium through regulation of E-cadherin which also affects the adjacent smooth muscle. Loss of E-cadherin led to apoptosis within Rab35 mutant kidney ureters resulting in widening of the ureter lumen. Our model is that Rab35 creates an Arf6 apical boundary through regulation of Arf6 activity, actin organization, through E-cadherin localization on the baso-lateral membrane. This regulation is necessary to maintain epithelial E-cadherin cell adherens and subsequently the tight junctions as well as membrane receptors that are associated with this response such as EGFR.

In summary, Rab35 mutant mice are a novel mouse model of non-obstructive hydronephrosis. We found that the role of Rab35 in regulating processes such as actin organization and primary cilia length is conserved across species and required for early embryonic development and tissue maintenance. Although we did not observe classic ciliopathic phenotypes at the timepoints assessed, this may be a result of the timing of developmental and juvenile inductions and allows for future investigation using tissue specific deletions. We showed that many of the mechanisms observed *in vitro* are recapitulated *in vivo* such as Rab35’s regulation of actin, E-cadherin, Arf6, and EGFR expression and localization. Finally, we showed that loss of Rab35 in kidneys and ureters leads to cell death. Our data provides valuable insight into not only the role of Rab35 *in vivo* but provides a novel avenue of study when assessing kidney and urinary system pathologies associated with hydronephrosis.

## Funding

This work was supported by National Institutes of Health [5R01HD089918-05 to JFR and BKY]; [2R01DK115751 to BKY]; [5T32GM008111-34 to KRC], [5T32DK116672-05 to KRC]

## Acknowledgements

We would like to thank members of Dr. Bradley K. Yoder’s laboratories for intellectual technical support on the project. We would like to thank Emily Helman and Dr. Jeremy Foote of the UAB Pathology Core for necropsy and pathology reports. We would like to thank the National Institute of Child Health and Human Development and the National Institute of Digestive and Kidney Diseases for funding these studies.

---

**Supplemental Figure 1.**
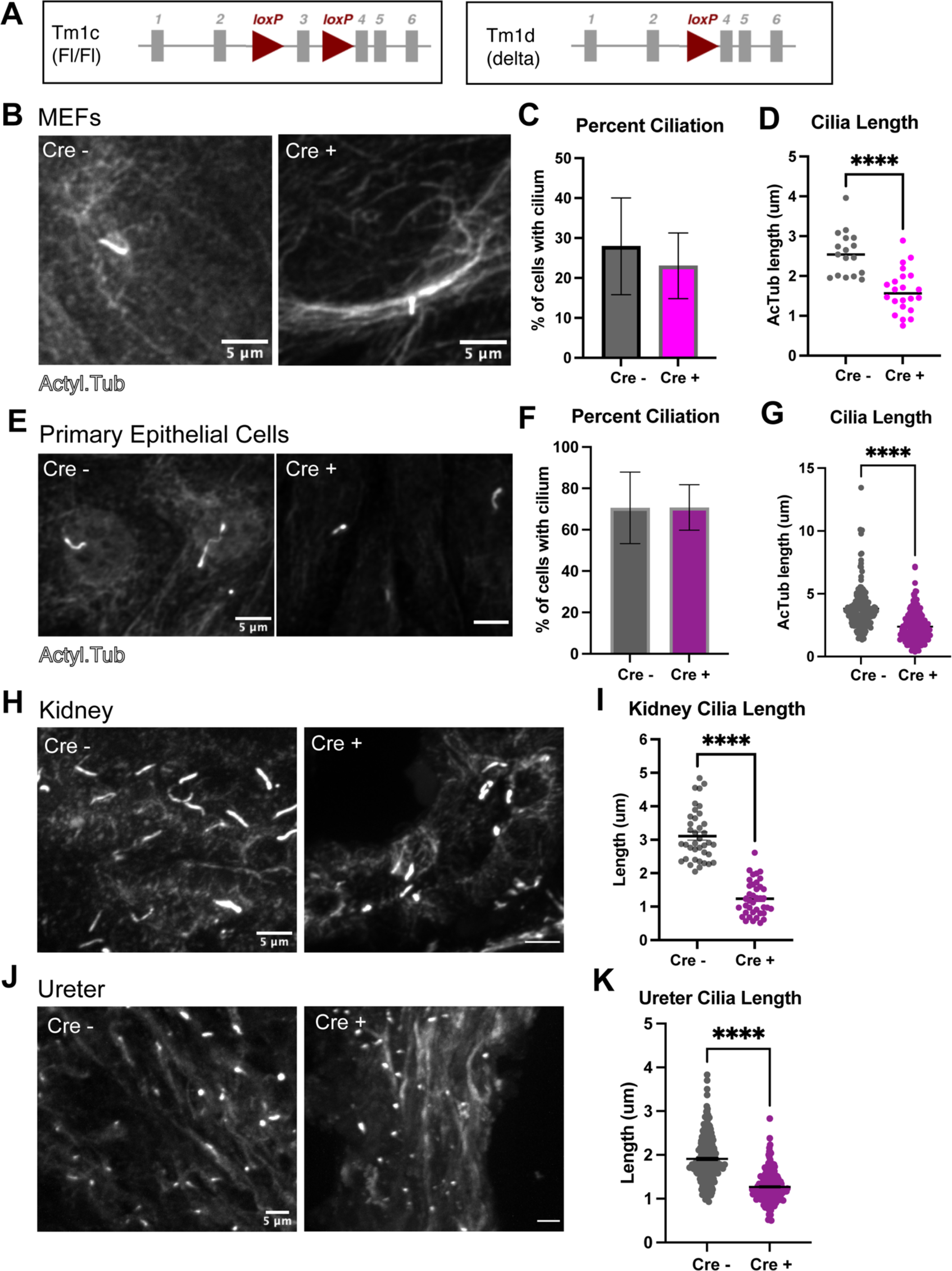
Conditional deletion of *Rab35* shortens cilia. **A)** Schematics of conditional (*tm1c*, *fl*) and deletion (*tm1d*, *delta*) alleles. **B)** MEFs isolated from E13.5 *Rab35^fl/fl^ Cre+* and *Rab35^fl/fl^ Cre-* embryos were treated with 4-hydroxytamoxifen in culture and stained for acetylated α-tubulin. We quantitated the percent ciliation **(C)** and cilia length **(D)**. **E)** Primary epithelial cells isolated from 1-month-old *Rab35^fl/fl^;Cre+* and *Rab35^fl/fl^; Cre-* mice, were treated with 4-hydroxytamoxifen in culture and analyzed similarly to MEFs **(F-G)**. **H)** Staining of *Rab35^delta^; Cre+* and *Rab35^fl/fl^; Cre-* control kidneys for acetylated α-tubulin. **I)** Quantitation of *Rab35^delta^; Cre+* and control kidney cilia length. **J)** Staining of *Rab35^delta^; Cre+* and *Rab35^fl/fl^;Cre-* control ureters for acetylated α-tubulin. **K)** Quantitation of *Rab35^delta^; Cre+* and control ureter cilia length. ****, unpaired two-tailed t test P value <0.0001.

**Supplemental Figure 2.**
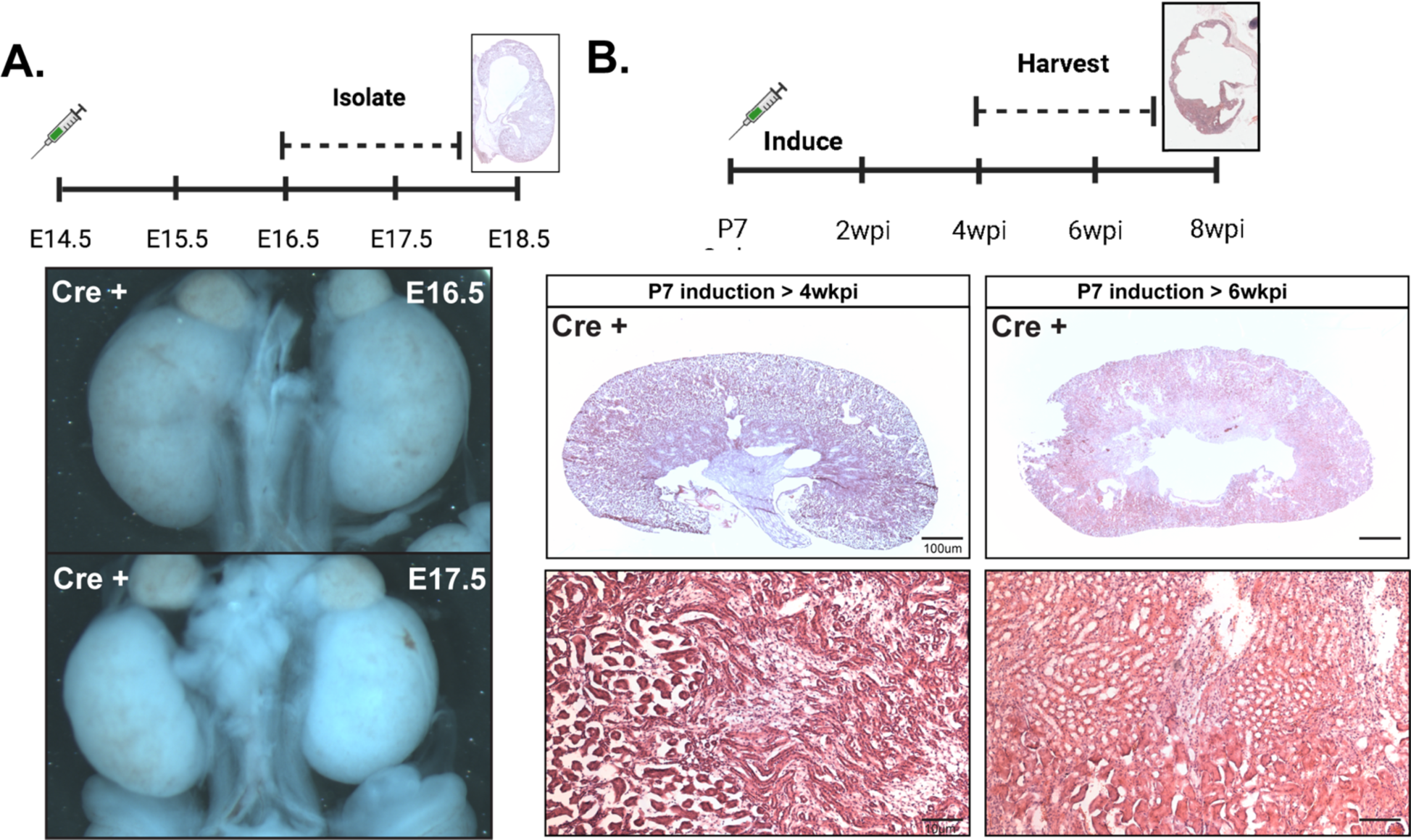
Progression of hydronephrosis during development and in the juvenile mouse. **A)** After E14.5 deletion, *Rab35^delta^;Cre+* renal histology at E16.5 and E17.5. Hydronephrosis developed after E16.5. **B)** After P7 deletion, *Rab35^delta^;Cre+* renal histology at 4 and 6 wkpi. Hydronephrosis developed after 4 wkpi. H&E staining of kidneys at 4wkpi and 6wkpi. Evidence of tubule dilation and hydronephrosis were first identified at 6wkpi.

**Supplemental Figure 3.**
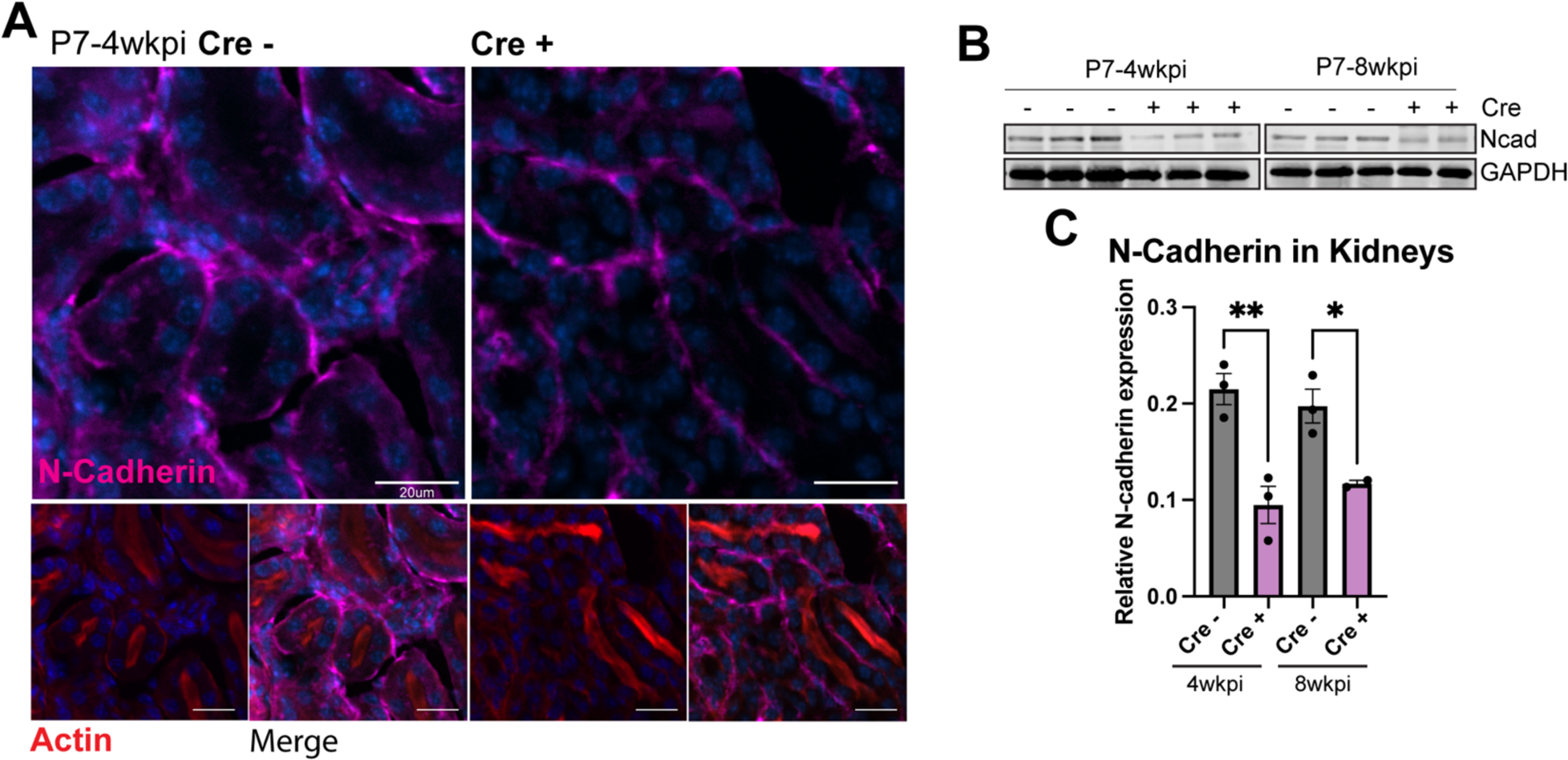
Expression of N-cadherin in kidneys. **A)** IF staining of control and pre-hydronephrotic kidneys to determine if Rab35 regulates other cadherins at the adherens junctions. There is no change in N-cadherin localization in pre-hydronephrotic kidneys but there are changes in expression **B)** by western blot of kidney lysates. Pre-hydronephrotic analysis P value= 0.0025 and hydronephrotic P value= 0.0340.

**Supplemental Figure 4.**
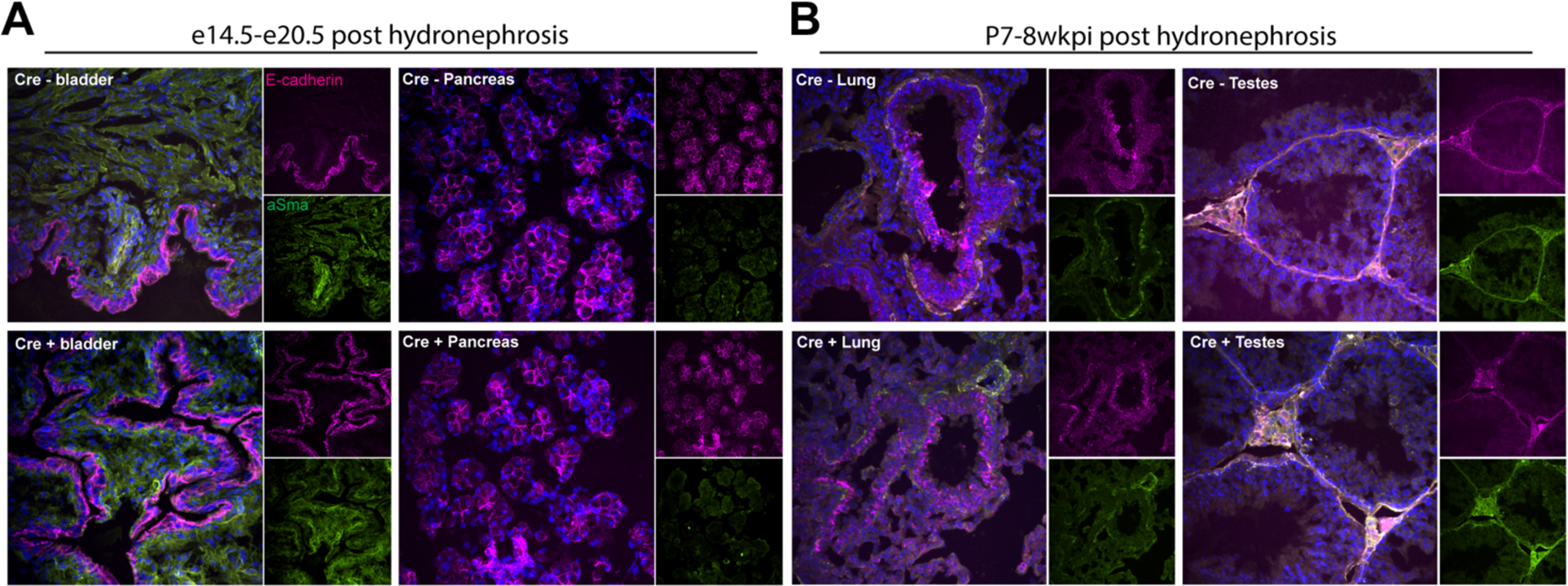
Loss of Rab35 does not affect other tissues at these timepoints. IF staining of tissues that may be affected by the loss of Rab35’s regulation of E-cadherin. E-cadherin and smooth muscle actin look comparable across **A)** bladder and pancreas in embryos with hydronephrosis compared to *Cre-* tissues, and **B)** the lung and testes of *Rab35^delta^;Cre+* animals with hydronephrosis compared to *Cre-* littermates.

**Supplemental Video 1.** Intrapelvic dye injections into control embryonic kidneys.

**Supplemental Video 2.** Intrapelvic dye injections into pre-hydronephrotic embryonic kidneys.

## Notes

### Competing Interest Statement

The authors have declared no competing interest.

